# The root of the transmissible cancer: first description of a widespread *Mytilus trossulus*-derived cancer lineage in *M. trossulus*

**DOI:** 10.1101/2020.12.25.424161

**Authors:** Maria Skazina, Nelly Odintsova, Maria Maiorova, Angelina Ivanova, Risto Väinölä, Petr Strelkov

**Affiliations:** Department of Applied Ecology, Saint-Petersburg State University, Saint-Petersburg 199178, Russia; A.V. Zhirmunsky National Scientific Center of Marine Biology, Far Eastern Branch of the Russian Academy of Sciences, Vladivostok 690041, Russia; Department of Ichthyology and Hydrobiology, Saint-Petersburg State University, Saint-Petersburg 199178, Russia; Finnish Museum of Natural History, University of Helsinki, Helsinki P. O. Box 17, FI-00014, Finland

## Abstract

Two lineages of bivalve transmissible neoplasia (BTN), BTN1 and BTN2, are known in blue mussels *Mytilus*. Both lineages derive from the Pacific mussel *M. trossulus* and are identified primarily by the unique genotypes of the nuclear gene EF1α. BTN1 is found in populations of *M. trossulus* from the Northeast Pacific, while BTN2 has been detected in populations of other *Mytilus* species worldwide but not in *M. trossulus* itself. The aim of our study was to examine mussels *M. trossulus* from the Sea of Japan (Northwest Pacific) for the presence of BTN. Using hemocytology and flow cytometry of the hemolymph, we confirmed disseminated neoplasia in our specimens. Cancerous mussels possessed the unique BTN2 EF1α genotype and two mitochondrial haplotypes with different recombinant control regions, similar to that of common BTN2 lineages. This is the first report of BTN2 in its original host species *M. trossulus* populations in West Pacific may be the birthplace of BTN2 and a natural reservoir where it is maintained and whence it spreads worldwide. A comparison of all available BTN and *M. trossulus* COI sequences suggests a common and recent, though presumably prehistoric origin of BTN2 diversity in populations of *M. trossulus* outside the Northeast Pacific.

## Introduction

Clonally transmissible cancer (CTC) is a neoplastic disease passed from individual to individual by physical transfer of cancer cells^1–3^. The first inkling of a transmissible cancer came from a study of canine transmissible venereal tumor, CTVT, dating back to 1876^4^. Since then CTC has been confirmed for CTVT^5^ and the facial tumor of Tasmanian devil *Sarcophilus harrisii*^6,7^ and, more recently, for several lineages of disseminated neoplasia (DN) of six marine bivalve mollusks: *Mya arenaria*^8^, *Cerastoderma edule*, *Polititapes aureus, Mytilus trossulus*^2^, *Mytilus edulis*^9^ and *Mytilus chilensis*^10^.

A straightforward method of CTC diagnostics is DNA genotyping. The genotype of CTC cancer cells is different from that of the host cells. The result is genetic chimerism, when an individual possesses cells with different genotypes^9^. At the same time, cancer cells of the same lineage have the same genotype in different infected individuals. Such lineage-specific genotypes are thought to derive from the “patient zero”, the host individual in which the cancer originated^2,8,11,12^.

The finding that CTC is the cause of DN in six different bivalve species is fairly recent, and it is reasonable to anticipate that further discoveries will turn up. There are also other lines of indirect evidence pointing to a widespread occurrence of transmissible cancers in bivalves^13^. At the same time, the data are not sufficient to ensure that CTC is the only cause of DN in those six species, or whether CTC is the usual cause of DN in bivalves in general.

DN is a fatal leukemia-like cancer affecting many marine bivalves^14^. It was first described in *M. trossulus* from the Northeast Pacific almost half a century ago^15^. DN is manifested in an uncontrolled proliferation of neoplastic cells in the hemolymph^16^, which ultimately replace healthy hemocytes. To note, though neoplastic cells are usually also referred to as hemocytes, their tissue origin remains unknown^14,17^.

Normal hemocytes of healthy bivalves (agranulocytes and granulocytes) are diploid and have a low mitotic activity^18,19^. Neoplastic hemocytes are aneuploid^18–22^, and have a high proliferation rate and a striking morphology. They are rounded, with a high nucleus-to-cytoplasm ratio^14,22,23^, and their nucleus is pleomorphic (i.e. variable in shape) and larger than that of normal hemocytes. As the disease progresses, the proliferating cells infiltrate other tissues, which leads to the loss of integration and normal functioning of organs^24,25^. DN may reach an epizootic level, causing mass mortality in wild and commercial bivalve populations^14,22,24^.

There are two approaches to DN diagnostics. The first approach is based on direct examination of cells in hemolymph smears or tissue sections. Various hemocytological techniques may be employed such as light microscopy of fresh or stained hemolymph smears^26,27^, microscopy of hemocytes with monoclonal antibodies specific to neoplastic cells^28^, and immunochemistry with antibodies specific to mitotic spindles (for revealing proliferating cells)^29^. The second approach is based on measuring DNA content. It employs flow cytometry of hemocytes labeled with fluorescent dyes specifically binding DNA^19^. Flow cytometry is less sensitive to the initial stages of DN than hemocytology but allows an efficient examination of large samples^19,22^.

Smooth-shelled blue mussels of the *Mytilus edulis* species complex are widely distributed in cold-temperate seas of both the Northern (*M. edulis*, *M. trossulus*, *M. galloprovincialis*) and the Southern (*M. galloprovincialis*, *M. chilensis*, *M. platensis* and *M. planulatus*) Hemispheres^30,31^ Mussels have been reported to harbor two presumably independent CTC lineages, BTN1 and BTN2^10^ (here BTN stands for bivalve transmissible neoplasia). Both lineages have been derived from *M. trossulus*, i.e. they have its genome. They are marked by different alleles of mitochondrial and nuclear loci, and the best single marker to distinguish the two lineages from each other and from hosts being the gene coding the elongation factor one alpha (EF1α)^2,10^. BTN1 was described in populations of its original host species in British Columbia (Northeast Pacific). BTN2 in turn has so far been found in *M. edulis* from Western Europe (Northeast Atlantic), in *M. chilensis* from Southern Chile (Southeast Pacific) and from the Beagle Channel in Argentina (Southwest Atlantic), but not in *M. trossulus* itself^2,10^. Yonemitsu *et al*.^10^ suggested that the original host, *M. trossulus*, could have evolved resistance to BTN2. Mitochondrial genomes of BTN2 clones from Chilean *M. chilensis* have been reported to be the product of recombination between the cancer (i.e. *M. trossulus-*derived) and the host (*M. chilensis*) mitogenome^10^.

So, *M. trossulus* gave rise to at least two CTCs, BTN1 and BTN2. BTN2 is a cross-species cancer, which is supposedly prone to occasional mitochondrial capture from the transient host and subsequent mitochondrial DNA recombination between the cancer and the host. Taken individually, these features are not unique. Two independent lineages of the facial tumor affecting Tasmanian devils are known^32^. Neoplastic hemocytes in the bivalve *P. aureus* have the genotype of another bivalve, *Venerupis corrugata*, for which CTC has not been reported^2^. Multiple episodes of mitochondrial capture from hosts and recombination between cancer and host mitogenomes are confirmed for CTVT^33^.

The total geographical distribution of BTN in the circum-global *M. edulis* complex remains poorly characterized, also as regards the original host species *M. trossulus*. *M. trossulus* is basically a North Pacific species, which then has spread to many areas of the North Atlantic in a series of repeated invasions^34–36^. In the Pacific it is distributed from the Arctic to the Gulf of California along the North American coast and to the Sea of Japan (hereinafter, SOJ) along the Asian coast^30^. So far only American Pacific populations of *M. trossulus* have been screened for BTN^2^. Riquet *et al*.^9^ and Yonemitsu *et al*.^10^ compared BTN mitochondrial sequences with all publically available *M. trossulus* sequences. Considering the common properties of sequences derived from BTN2, from a mussel individual 62mc10 of *M. trossulus* from the Baltic Sea (NCBI accession number KM192133^37^) and individuals from Norway^38^, Yonemitsu *et al*.^10^ hypothesized that the BTN2 lineage originated in a North European *M. trossulus* population, i.e. outside of the ancestral range of this species. DN has indeed been recorded in *M. trossulus* in the Baltic Sea^39^. However, DN was also found in the SOJ^40^, within the ancestral range of *M. trossulus* where the presence of BTN has so far not been assessed. In the Northeast Pacific, on the North American coast, mussels are infected with BTN1, but the possible occurrence and identity of the infections on the opposite side of the Ocean are unknown.

The aim of our study was to examine the populations of *M. trossulus* from the SOJ for the presence of BTN and, should it be found, to establish its genetic affinity with the known BTN lineages.

## Material and methods

### Study design

The workflow of the study is depicted in Figure 1. In brief, mussels were collected in the Gulf of Peter the Great, the SOJ. Different tissues were sampled from each individual (“*Sample collection and preprocessing”* step). The hemolymph of all mussels was analyzed by flow cytometry. Based on the results, the mussels were preliminary classified as cancerous (DN-suggested) if an increased fraction of aneuploid hemocytes was registered. For five mussels (four DN-suggested and a healthy control) the diagnosis was verified by direct examination of hemocytes using hemocytology and immunochemistry (“*DN diagnostics*”). The same cancerous mussels and the controls were used during the next steps. There, we generally followed the methodology of previous BTN studies^2,9,10^. Hemolymph and foot tissues were genotyped separately for nuclear microsatellites and EF1α, and for the mitochondrial COI and control region (CR). For EF1α and CR, when multiple alleles were present and could not be resolved by sequencing, molecular cloning was employed. Two kinds of evidence for the presence of BTN were considered: genetic chimerism of cancerous individuals, with the hemolymph and the foot tissues being dominated by different sets of alleles, and the identity of “extra” alleles of different cancerous individuals (see Introduction) (“*BTN diagnostics*”). At the step of “*Phylogenetic analyses*”, evolutionary relationships of cancer-associated genotypes were assessed. Firstly, all relevant BTN sequences from the previous studies were considered in order to identify the known cancer lineages, if any, affecting mussels from the SOJ. Secondly, all available *M. trossulus* COI sequences were examined in order to find out whether the cancer alleles identified in the SOJ mussels had been recorded anywhere before and whether they demonstrate an affinity to particular mitochondrial lineages of the host. Finally, we verified the purebred *M. trossulus* ancestry of mussels from BTN-infected population by genotyping them by three additional taxonomically diagnostic markers (“*Species confirmation*”) and identified their sex histologically and/or genetically (*“Sex identification”*).

**Figure 1.**
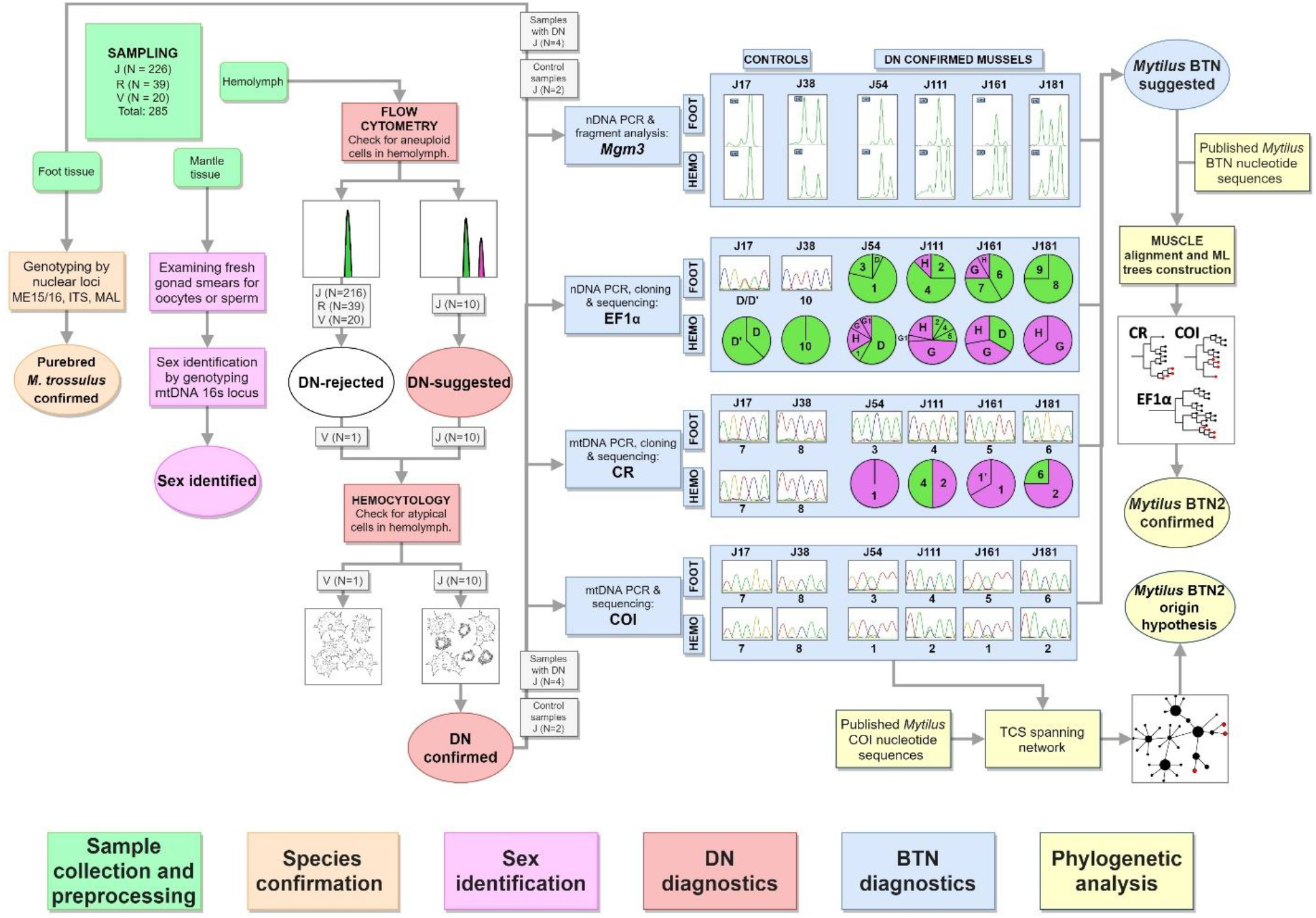
Study design. Different steps of work are color-coded (see boxes at the bottom). Unique numbers of mussels used at different steps are given. Main results of the analyses are schematically outlined (see text). Real microsatellite electropherograms of *Mg*μ3 are shown. Sectors of pie charts are frequencies of sequences revealed by molecular cloning (sequences found in one or just a few colonies are not considered). Sequences are indicated by their unique names; supposedly cancerous alleles are shown in purple.

### Sample collection and preprocessing

Mussels were collected by scuba diving at three localities of the SOJ in July-September 2019: Vladivostok city public beach “Vtoraya Rechka” (43°10’02”N, 131°58’07”E, depth 5 m, a natural bottom habitat, sample size N=39, mean shell size L=25 mm, sample “R”), “Vostok” Marine Biological Station (42°89’35”N, 132°73’33”E, depth 5 m, experimental mussel plantation, N=20, L=50 mm, sample “V”) and the Gaydamak Bay (42°52’04”N, 132°41’27”E, fouling of a mooring buoy anchored at a depth about 5 m about 50 m from the shore, N=226, L=37 mm, sample “J”). “Vostok” station is a nature reserve and the least polluted of the studied localities, while Gaydamak is an industrial harbor heavily polluted by sewage. After sampling, the mussels were transported to the laboratory and stored for 2-4 days alive, each sample in a separate aquarium, before the experiments.

Hemolymph was collected from the posterior adductor with the help of a syringe with a 22-gauge needle. Some aliquots of hemolymph were used immediately for flow cytometry, and some were fixed in 4% PFA for future immunochemistry studies. Some aliquots, as well as pieces of mantle and foot tissues from each specimen, were fixed in 70% ethanol for genetic analyses.

### DN diagnostics

#### Flow cytometry

Flow cytometry analysis and the interpretation of the results followed Vassilenko & Baldwin^19^. Hemocytes stained with 4’-6-diamidino-2-phenylindole (DAPI) were analyzed on a CytoFLEX flow cytometer with CytExpert software (Beckman-Coulter, USA). Plots of side scatter SSC-A versus forward scatter FSC-A were used for visualization of cell groups. Duplets were removed by gating on forward scatter height FSC-H against forward scatter FSC-A. Histograms of fluorescence signal PB450-A were used for determination and gating of cells with different levels of ploidy. At least 10,000 events were evaluated for each sample. Histograms of PB450-A were used to estimate ploidy levels of aneuploid peaks relative to the diploid peak of the same specimen and the rate of aneuploid cells in the sample. Mussels were diagnosed as healthy (“DN-rejected”) if the scatter plot SSC-A *vs* FSC-A and PB450-A histogram of fluorescence indicated only non-proliferating cells (agranulocytes and granulocytes) with one peak for diploid phase or with admixture of very few (<5%) proliferating cells (minor peak for tetraploid cells). If a cell population with additional peaks was detected, the individuals were considered as “DN-suggested”.

#### Hemocytology and immunochemistry

All procedures for cell fixation and staining were described in detail in a previous study^41^. In brief, the cells were stained with TRITC-labeled phalloidin (Molecular Probes) and DAPI (Vector Laboratories) for visualization of actin cytoskeleton and nucleus. We used primary mouse monoclonal antibodies against anti-α-acetylated tubulin (clone 6-11B-1, Sigma Aldrich and proliferating cell nuclear antigen (PCNA, clone PC10, Abcam) for detecting mitotic spindles and proliferating cells, respectively. These antibodies were previously characterized as labeling dividing cells in bivalves^41–44^. Observations and images were made with Zeiss LSM 780 confocal laser scanning microscope. A mussel was considered as “DN-confirmed” if non-adherent hemocytes with a high nucleus-to-cytoplasm ratio and a pleomorphic nucleus were detected.

### BTN diagnostics

Total DNA was extracted separately from the hemolymph and the foot tissues using the DNeasy Blood and Tissue Kit (Qiagen, USA) according to the manufacturer’s protocol.

The microsatellite marker *Mg*μ3, polymorphic in *M. trossulus,* was amplified according to the protocol in the original study^45^ and subjected to fragment analysis by capillary electrophoresis. Microsatellite fragment patterns were analyzed in GelQuest (https://www.sequentix.de/gelquest/help/index.html). The attempt to amplify other microsatellite loci (*Mg*μ5, *Mg*μ6, *Mg*μ7) was not successful.

Three DNA fragments—an intron-spanning region of the nuclear EF1α gene, a fragment of the mitochondrial COI and a fragment spanning 16S rRNA and CR—were amplified using primer sequences and PCR cycling conditions as in Metzger *et al*.^2^ and Yonemitsu *et al*.^10^ and sequenced in both directions by the Sanger method. The primers used for COI amplification target the female mitochondrial genome (F-mtDNA), while primers for 16S and CR potentially amplify both male (M-mtDNA) and F-mtDNA^10^. Due to the doubly uniparental inheritance (DUI) blue mussel females are homoplasmic for F-mtDNA, which is inherited maternally, while males are heteroplasmic, carrying the F-mtDNA in somatic tissues and the paternally inherited M-mtDNA in gonads^46^.

Sequencing and fragment analysis were performed on Genetic Analyzer ABI Prism 3500xl at the Centre for Molecular and Cell Technologies of St. Petersburg State University Research Park (https://researchpark.spbu.ru/en/biomed-eng). Sequence chromatograms were visually studied in MEGA Х software^47^.

EF1α PCR products from the hemolymph and the foot tissues of the cancerous and the control mussels were subjected to molecular cloning. CR PCR products were cloned from the hemolymph of the cancerous mussels only. Molecular cloning procedures were subcontracted to Evrogen JSC (Russia). Quick-TA kit (Evrogen JSC) was used for cloning, and the plasmids were transformed into competent *E. coli* (Evrogen JSC). In all the cases at least 16 colonies were sequenced using M13 primers. Some sequences were detected only in one colony. They were probably artificial mutations generated by PCR and molecular cloning procedures such as polymerase errors and random crossing-over of incomplete PCR extension products of original alleles^48,49^ and therefore were excluded from the following analyses.

In COI, signals from multiple alleles were resolved by sequencing only. If “piggybacks”, that is, overlapping peaks at some positions, were observed on chromatograms, the major peaks were attributed to the presumable cancer allele in the hemolymph samples and to the presumable host allele in the foot samples.

Sequence chromatograms were analyzed in MEGA X^47^. Sequences were aligned with MUSCLE algorithm with some manual adjustment.

#### Phylogenetic analysis

Nucleotide sequences of CR, COI and EF1α from four cancerous mussels were aligned together with the corresponding sequences from a previous BTN study (data on 11 mussels with confirmed BTN)^10^ and mitochondrial sequences of the Baltic mussel 62mc10^37^. The 62mc10 genome was previously shown to be similar to cancerous ones^10^. Alignments by MUSCLE and maximum likelihood phylogenetic trees were generated using MEGA X^47^, with 100 bootstrap replicates, treating gaps in the alignment as missing data. The same substitution models as in Yonemitsu *et al*.^10^ were used for tree generation. The trees were visualized using iTOL tool^50^. The analysis of BTN2 CR alleles in Yonemitsu *et al*.^10^ revealed that they are recombinant molecules with the insertion of M-mtDNA segment into the F-mtDNA. Therefore we employed the RDP4 package^51^ to detect possible recombination breakpoints in the cancer-associated CR alleles found in our study (Supplementary Fig. S5).

In the comparative analyses of *M. trossulus* and BTN COI sequences, we considered the data from this study (four cancerous and 20 healthy mussels), those from previous BTN studies summarized by Yonemitsu *et al*.^10^ and all publically available *M. trossulus* COI data^10,36,37,52–55,56,57^. The alignment was created by ClustalW algorithm in MEGA X^47^. COI haplotype diversity was visualized using the TCS haplotype network^58^ built within the PopART software^59^. *M. trossulus* samples were classified by their geographical origin into four groups (“macroregions”), i.e. the Northwest Pacific, Northeast Pacific, Northwest Atlantic and Northeast Atlantic.

#### Species confirmation and sex identification

According to a recent survey^60^, the blue mussel populations in the SOJ are overwhelmingly dominated by *M. trossulus*, but the presence of a “cryptic” species, *M. galloprovincialis* and its hybrids with *M. trossulus* cannot be entirely ruled out. Mussels from the “J” sample (N=21), including four target mussels with DN, were genotyped for three nuclear markers routinely used for discriminating *M. trossulus* and *M. galloprovincialis*^61^, ME15/16^62^, ITS^63^ and MAL-I^64^, using the primers and protocols in the original articles. The DNA extracted from the hemolymph and from the mantle of each mussel were analyzed in parallel. The mussels were also sexed, at first by a microscopic examination of fresh tissues of the mantle, where the gonads in blue mussels are partly localized. It turned out, however, that most mussels were in post-spawning condition and lacked gametes. Therefore we identified their “mitochondrial” sex by the presence or the absence of M-mtDNA 16S haplotypes, following the approach of Rawson and Hilbish^65^ and using DNA extracted from the mantle.

## Results

### DN diagnostics

#### Flow cytometry

Flow cytometry of hemolymph revealed two distinct patterns (Fig. 2). Most of the individuals had one peak of hemocytes, interpreted as diploid (Fig. 2a), or a diploid peak with a small admixture of tetraploids (Fig. 2b). These mussels were diagnosed as healthy. The second pattern was revealed in 10 “J” individuals (4% of “J” sample), which had an additional population of aneuploid cells (Fig. 2c-f). Their ploidy, calculated relatively to the 2n peak, varied among individuals from 3.7n to 5.2n. The proportion of these cells in the hemolymph of different mussels varied from 12 to 98%. These mussels were classified as “DN-suggested”. We did not reveal any significant signal from proliferating neoplastic cells, which could be expected as a third peak at the right side of the aneuploid peak in the histogram. Two “DN suggested” mussels with a moderate proportion of neoplastic hemocytes, J54 (proportion 44.4%) and J111 (26.4%), and two mussels with a high proportion of neoplastic hemocytes, J161 (91%) and J181 (80%) (Fig. 2c-f), were chosen for further analyses along with the control V1.

**Figure 2.**
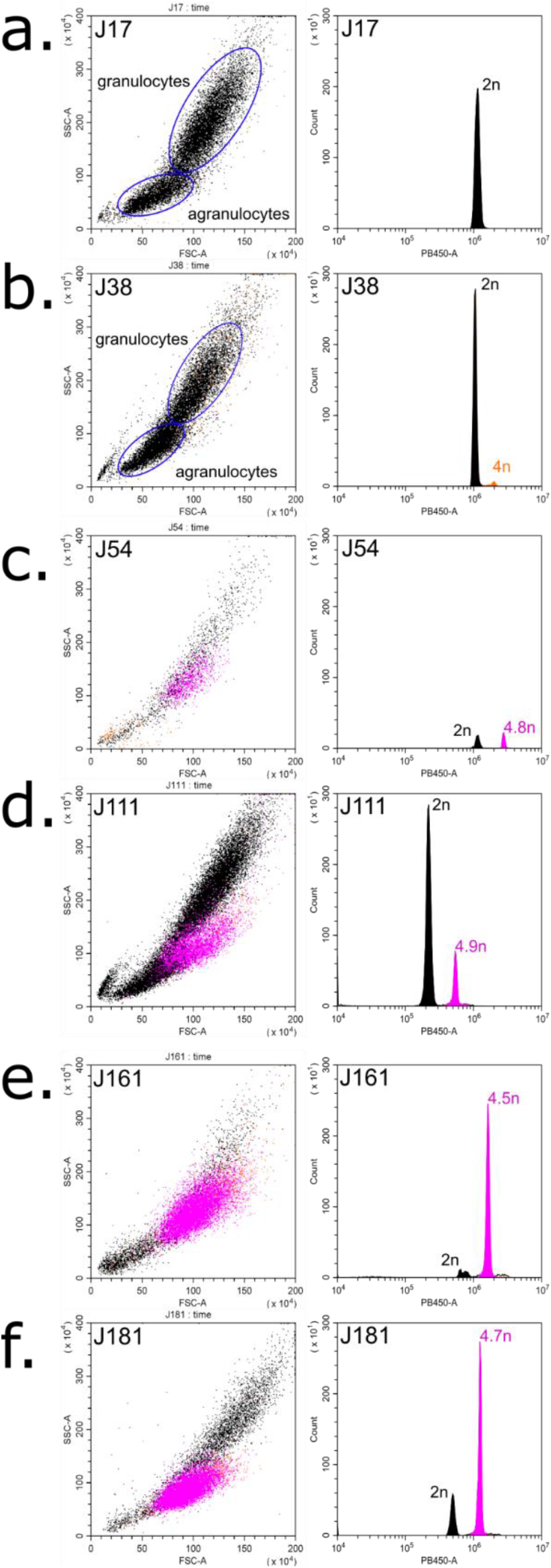
DN diagnostics in individual *M. trossulus* mussels from the Gaydamak Bay, the SOJ, by flow cytometry. Flow cytometry plots depict cell groups (left graphs) and histograms revealing ploidy levels (right graphs) for healthy (**a, b**) and DN-suggested (**c-f**) mussels. An additional population of aneuploid cells (violet peaks and dots) is observed in the hemolymph of each DN-suggested mussel. Normal non-proliferating hemocytes are shown in black, normal proliferating hemocytes (tetraploid), in orange. Relative ploidy levels are given near the peaks. Subpopulations of agranulocytes and granulocytes are highlighted for healthy mussels (**a, b**).

#### Hemocytology and immunochemistry

Hemocytological study revealed striking differences between the control mussels on the one hand and all the four DN-suggested mussels on the other hand. DN-suggested individuals had, in addition to normal adherent hemocytes with a low nucleus-to-cytoplasm ratio, also anomalous round non-adherent hemocytes. They looked like hedgehogs on the slides, the resemblance being due to an altered cytoskeleton with prominent actin “spikes”. Their nuclei were pleomorphic, larger than those of normal hemocytes (Fig. 3). We considered these cells as neoplastic and the mussels they belonged to as DN-confirmed.

**Figure 3.**
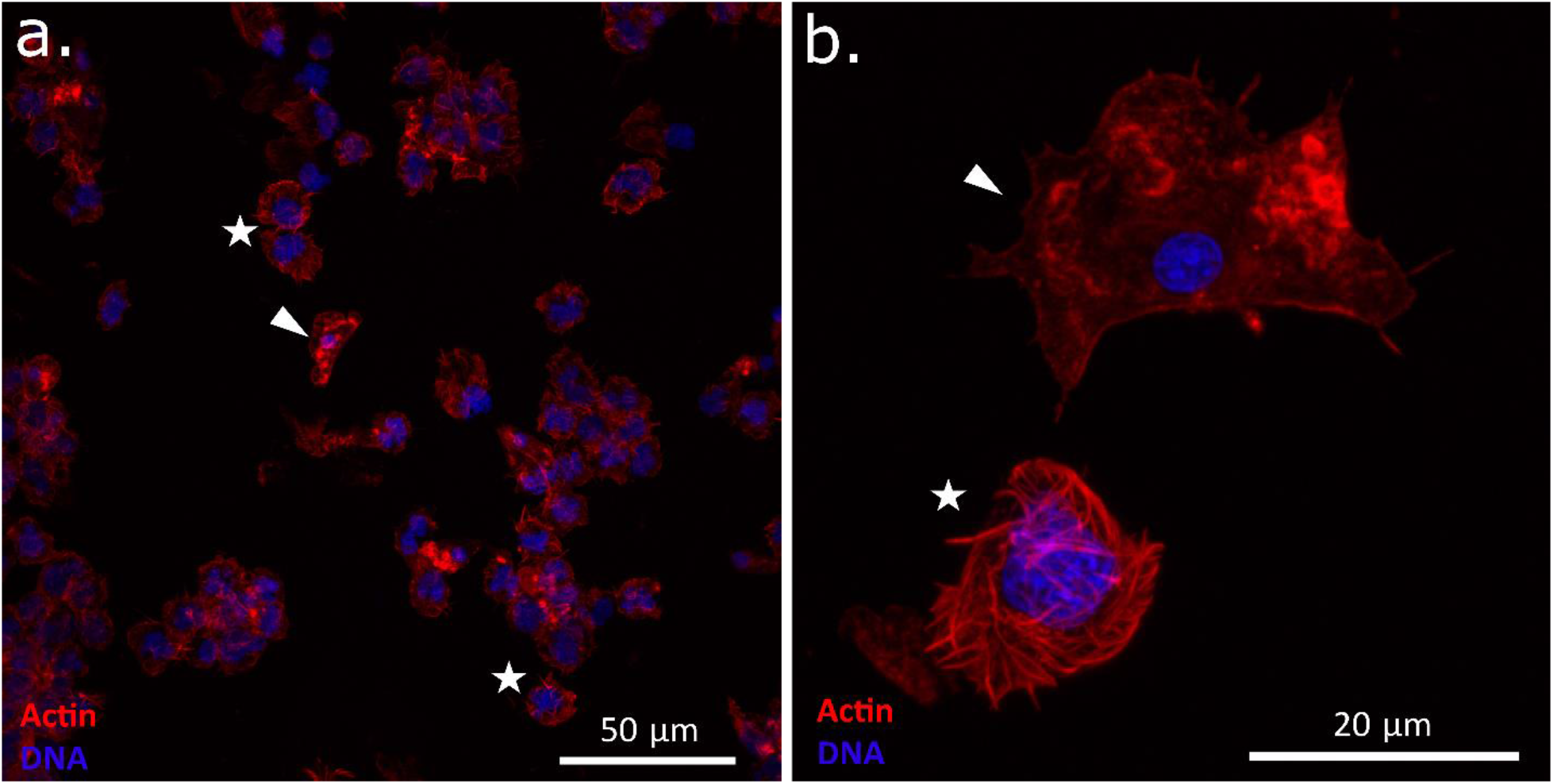
Confocal microscopy images of hemocytes of a DN-suggested mussel (J54) stained with DAPI (blue) and TRITC-labeled phalloidin (red). Images (**a**, **b**) are at two different magnifications (note the scale bars). Arrowheads point to normal adherent hemocytes with a small compact nucleus. Stars mark neoplastic aneuploid round cells with a large lobed nucleus and an altered actin cytoskeleton.

PCNA staining revealed few neoplastic hemocytes at the DNA synthesis stage (Supplementary Fig. S1). This observation was in line with the results of flow cytometry, which showed a low proliferating rate of the neoplastic cells.

### BTN diagnostics

In the microsatellite analysis the fragment size of the *Mg*μ3 marker varied in the range of 141-146 bp. In healthy specimens (J2-J17, J38) the size and the representation of the amplified fragments (position and height of the peaks on the electropherograms, Supplementary Fig. S2, see also Fig. 1) from the hemolymph and from the foot samples were the same. In contrast, in all the four DN-confirmed mussels the 146 bp fragment was overrepresented in the hemolymph samples in comparison with the foot samples (Fig. 1, Supplementary Fig. S2). At the same time, this fragment, as well as the 144 bp fragment, characteristic of all the cancerous mussels, was also recorded in some healthy mussels, making the results inconclusive.

In sequenced marker genes, the lengths of the fragments in the final alignment were 447-453 bp for EF1α, 622-624 bp for CR and 630 bp for COI. The cloning results are summarized in Supplementary Table S2. The NCBI GenBank accession numbers are in Supplementary Table S3. Hereafter the alleles identified in our study will be designated by letters if initially recorded in the study of Yonemitsu *et al*.^10^, and by numerals if newly found, except in a few cases when we wanted to emphasize the similarity between the alleles (e.g. EF1α-G1, CR-1’, see below). It should be noted, however, that the alignments in Yonemitsu *et al*.^10^ were slightly longer than here.

The sequencing and molecular cloning of EF1α PCR products revealed no differences between sequences from the hemolymph and from the foot tissues of the control individuals (Supplementary Table S2 and Fig. 1). In contrast, all EF1α sequencing chromatograms from the hemolymph and the foot samples of DN-confirmed mussels showed a mixed signal. Molecular cloning of these PCR products revealed 2-4 different sequences in the foot samples and 2-6 in the hemolymph samples (Fig. 1, Supplementary Fig. S3). Some sequences were common, that is, represented in relatively many colonies, and differed by more than one substitution, while others were rare. All the mussels had two common sequences, EF1α-G and EF1α-H. They differed by 21 substitutions, which made them the most dissimilar common sequences, and were always more frequent in the hemolymph than in the foot samples. Other common sequences were usually specific of individual mussels and were present in both tissues (Fig. 1, Supplementary Table S2 and Supplementary Fig. S3). Putatively, EF1α-G and EF1α-H represented a heterozygous cancer genotype. Other common sequences probably represented the diploid host genomes. Rare sequences, including EF1α-G1 different from EF1α-G by one substitution (Supplementary Fig. S3), were probably methodological artifacts.

Direct sequencing of COI and CR revealed identical alleles in the hemolymph and in the foot samples of individual control mussels, in homoplasmic condition (J17 and J38, Supplementary Table S2). On the contrary, the chromatograms from analyses of different tissues of mussels with DN looked very different.

No heteroplasmy was observed in the CR chromatograms of cancerous mussels. In foot samples, unique alleles were identified. In the hemolymph samples, two alleles were identified: CR-1 in J54 and J161 and CR-2 in J111 and J181. CR-1 and CR-2 were very different both from each other (31 substitutions, 5.0% difference) and from the other alleles (2.7-6.6%), considering that the differences between all the other alleles were within the range of 0.3-1.5%. Molecular cloning confirmed the results of direct sequencing, but also revealed additional rarer sequences invisible on sequencing chromatograms (Supplementary Table S2, Fig. 1). In the hemolymph of J111 and J181 the same alleles as in foot were found, while in the hemolymph of J161 an allele CR-1’, supported by five clones, was found, differing from the major CR-1 allele by four substitutions.

No COI heteroplasmy was observed in the foot samples of cancerous mussels while several positions with overlapping peaks of a very different height were observed on chromatograms of the hemolymph samples (cf. Fig. 1). The heteroplasmy was readily identified as representing presence of the “foot allele” of the same individual in minority (lower peaks) in addition to a dominant hemolymph allele. Two major COI sequences were identified in the hemolymph samples: COI-1 in J54 and J161 and COI-2 in J111 and J181. These alleles differed from each other by 6 substitutions (0.95%) and by 0.60-0.95% from all the other alleles.

Thus, the sequence analyses revealed genetic chimerism of mussels with DN, with the hemolymph and the foot tissues being dominated by different genotypes of both the nucleus and mitochondrion. However, while EF1α genotyping revealed the same cancer genotype in all the diseased mussels, mtDNA genotyping revealed two different cancer genotypes, marked by different combinations of COI and CR alleles, in different mussels. The conclusive evidence that DN in mussels from the SOJ is BTN came from a genetic comparison with cancers from previous studies.

#### Phylogenetic analysis

##### Maximum likelihood trees

A general inspection of phylogenetic trees (Fig. 4) shows that the SOJ mussels are infected with the BTN2 cancer lineage. The EF1α-G and EF1α-H alleles identified in these mussels are previously known BTN2-specific alleles from other *Mytilus* hosts. For both mtDNA fragments the cancerous alleles clustered together with the major BTN2-specific alleles and the 62mc10 (Fig. 4). The alleles COI-1 and COI-2 differed from the major COI-B BTN2 allele by 6 and 8 substitutions, respectively. CR-1 allele differed from the closest CR-D allele by 16 substitutions, and from the 62mc10 by 11 substitutions. CR-2 differed from the closest CR-C allele by 5 substitutions. The alleles of *M. trossulus* (i.e. host alleles) from the SOJ were randomly scattered across the *M. trossulus* clade on the trees (Fig. 4).

**Figure 4.**
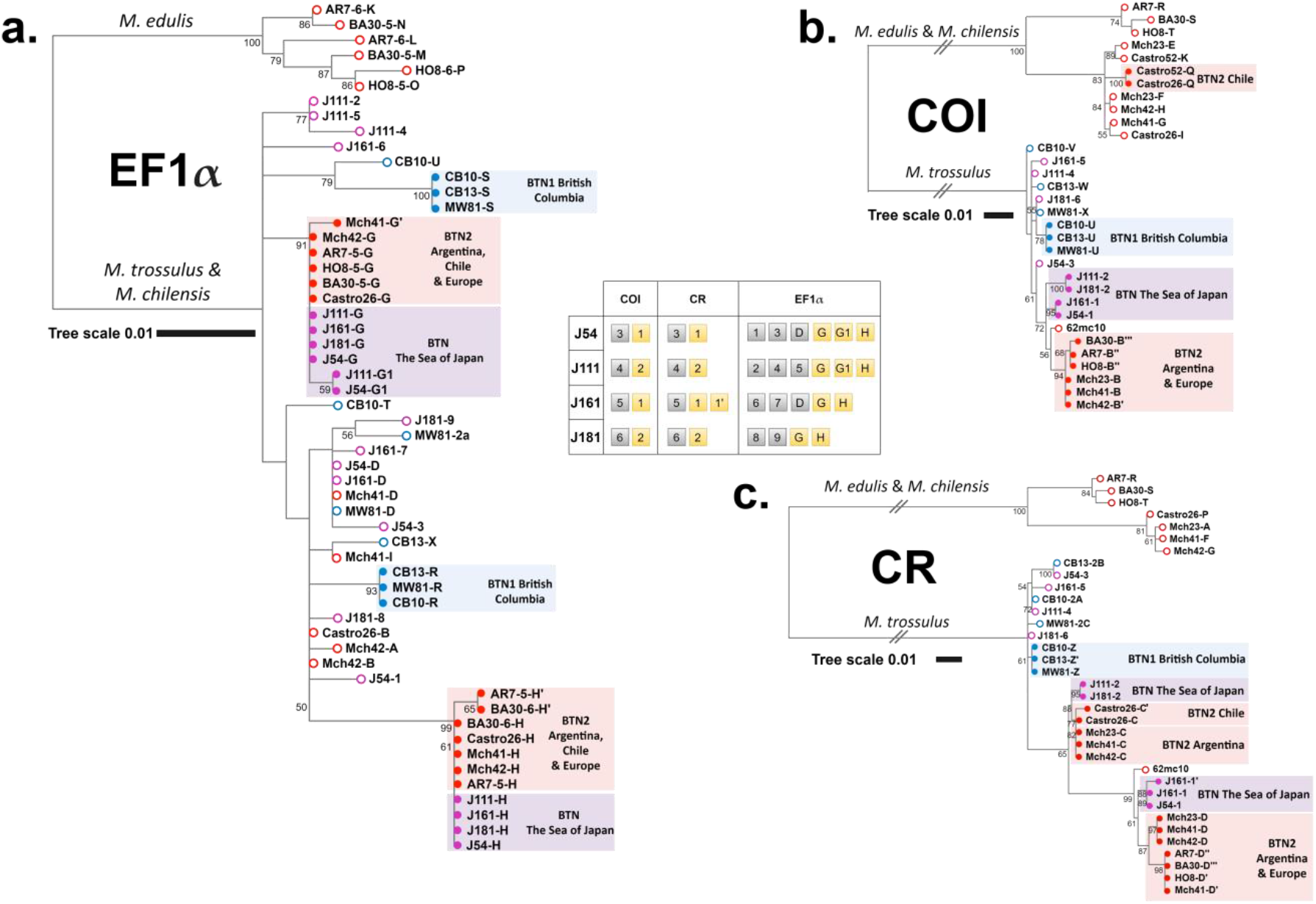
Phylogenetic analysis of nuclear and mitochondrial loci from mussels with BTN. (**a**) EF1α maximum likelihood tree based on the 629 bp alignment (HKY85+G substitution model) (**b**) COI maximum likelihood tree based on the 630 bp alignment (GTR+G substitution model) (**c**) CR maximum likelihood tree based on the 842 bp alignment (HKY+G substitution model). Maximum likelihood trees were rooted at the midpoint, with bootstrap values below 50 removed. Branches marked by two slashes were shortened 5-fold for COI and 3-fold for CR. The scale bars mark genetic distance. Source data include all references from Yonemitsu *et al*.^10^ with original allele nomenclature (specimen ID, allele ID; Castro26 host COI allele J not included because of the short length of sequence), mitochondrial sequence KM192133 from the Baltic mussel 62mc10 and new data on four mussels with DN from the SOJ. Open circles at the end of branches mark host alleles, closed circles mark cancer alleles. Alleles revealed in mussels with BTN1 are marked in blue, those revealed in mussels with BTN2, in red. Alleles revealed in mussels from the SOJ are marked in violet (alleles are listed in the legend). Geographic origin of the samples and species identity of clades are indicated. Alignments are provided in Supplementary data S1-S3.

An analysis of recombination identified the same breakpoints in CR-1/CR-1’ and CR-D alleles but different breakpoints in CR-2 and CR-C alleles (Supplementary Fig. S5). However, CR-2 and CR-C differed by only two substitutions between the suggested breakpoints. We will therefore adhere to the hypothesis that CR-1 and CR-D, as well as CR-2 and CR-C, represent the same mitochondrial lineages originated through singular recombination events. To remember, individual J161 was heteroplasmic for the close CR-1 and CR-1’ alleles. Such “additional” heteroplasmy occasionally occurs in BTN2 in different parts of its geographical distribution (Mch-41, Castro-26, Fig. 5).

**Figure 5.**
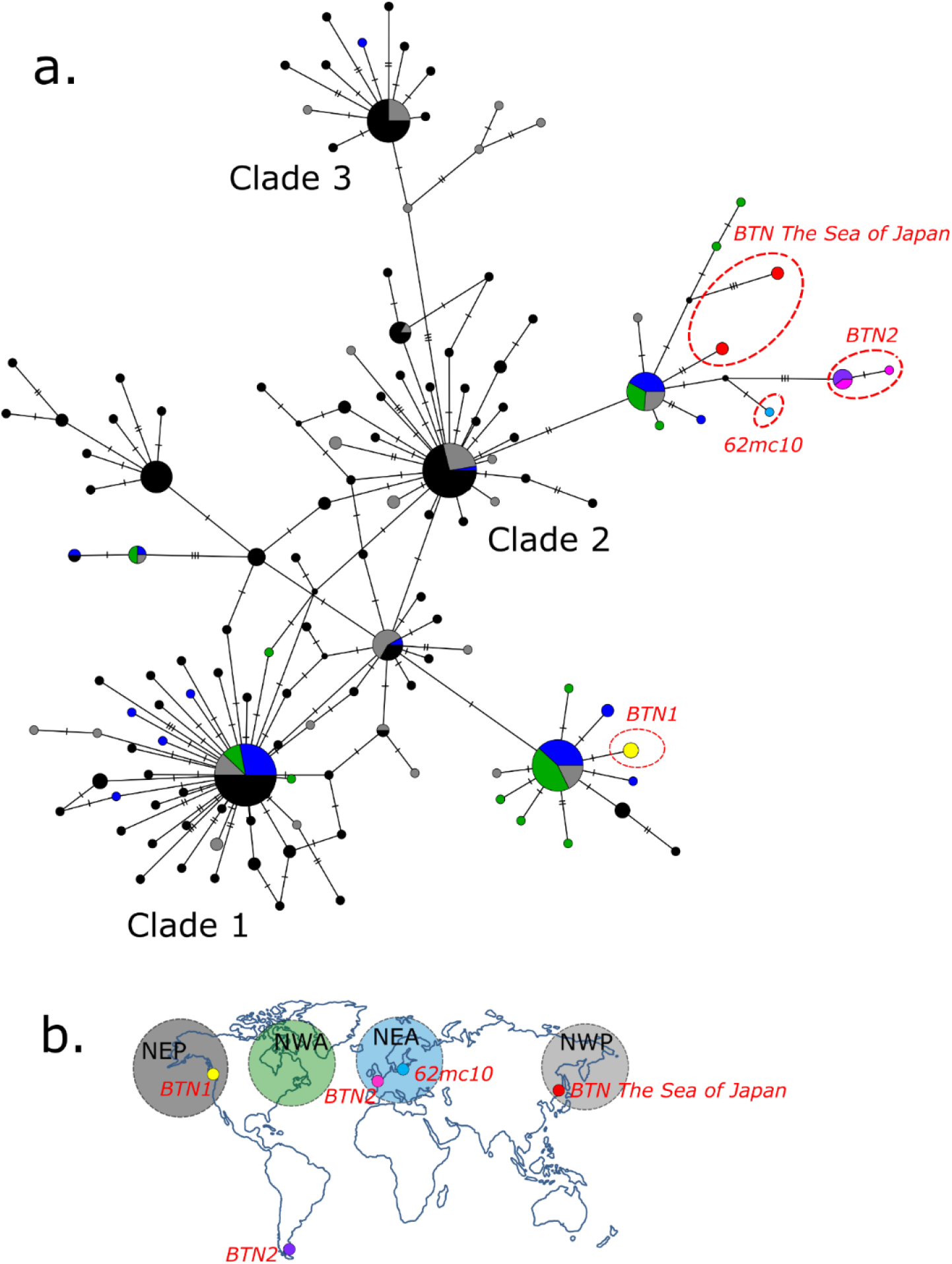
(**a**) *M. trossulus* and BTN COI haplotype network obtained from the TCS analysis. The analysis was based on the 542 bp alignment (provided as Supplementary data S4) of 350 sequences produced in this study (4 cancerous and 20 healthy mussels) and from previous BTN studies, together with all publically available *M. trossulus* sequences (except the dataset from Vancouver Island, British Columbia^54^, see the list of sequences in Supplementary Table S1). Each circle represents a single allele. The size of the circle is proportional to the number of individuals bearing the allele. Bars indicate mutations between alleles. Small black circles indicate hypothetical haplotypes predicted by the model. The geographical origin of samples is color-coded and illustrated in the map (**b**). Samples corresponding to BTN1 (all from British Columbia) are in yellow, to BTN2 in pink (Europe) and purple (Argentina), to BTN from the SOJ in red. Reference *M. trossulus* samples from the Northwest Pacific (NWP on the map) are in grey, from Northeast Pacific (NEP), in black, from Northwest Atlantic (NWA), in green, from Northeast Atlantic (NEA), in dark blue, and from the Baltic Sea (a single sample 62mc10), in light blue. Clades 1-3 are named after Marko *et al*.^53^. See Supplementary Fig. S4 for the results of a re-analysis of these data together with those from Crego-Prieto *et al*.^54^

In addition to the genetic similarity of BTN2 worldwide, Fig. 4 also illustrates geographic differences within this cancer. In the SOJ there are basically two cancer mtDNA haplotypes, comprising two 1% divergent COI alleles and two very divergent CR alleles (CR-1, CR-2), while elsewhere there are two cancer haplotypes, comprising basically the same COI alleles of the “B group” and two CR alleles (CR-C, CR-D) similar to that in the SOJ. Another difference is that the two cancer haplotypes are apparently homoplasmic in the SOJ and in Europe but heteroplasmic in Argentina (both alleles present in the same cancerous mussels).

##### Haplotype network

Among the COI data sets included in the analysis, that from Vancouver Island, British Columbia^54^ demonstrated a remarkably strong polymorphism in comparison with other regional sets, including another Northeast Pacific set. Therefore we provide separate networks performed without and with the data of Crego-Prieto *et al*.^54^ in Fig. 5 and Supplementary Fig. S4, respectively. The analyses revealed a complex star-like haplotype network familiar from the previous phylogeographic studies^34,36,53^. The network consisted of several major clades, each with a common core haplotype and many rare haplotypes radiating from it. Some clades were “cosmopolitan” and included samples from both the Atlantic and the Pacific oceans, while others were nearly restricted to the Pacific. The unique BTN1 allele belonged to one of the cosmopolitan clades, which included sequences from all the four macroregions considered but was dominated by samples from the Northeast Pacific. The BTN2 alleles, including that from the SOJ, and the Baltic 62mc10, were attached to another clade. This clade included samples from all the macroregions except the Northeast Pacific (Fig. 5), with one notable exception. A single sequence identical to the cancerous COI-1 from the SOJ was identified in the data from Bamfield locality on Vancouver Island (KF931805^54^, Supplementary Fig. S4).

#### Species confirmation and sex identification

Only *M. trossulus* alleles were recorded in a subsample of mussels (N=21, including J17, J38, and cancerous J54, J111, J161 and J181) studied by three additional nuclear markers, and the genotypes retrieved from different tissues of the same mussels were always the same. Twelve mussels were identified as “mitochondrial males” by 16S locus, among them J181, and the other mussels, as females.

## Discussion

In this study we showed that *Mytilus trossulus* mussels from the SOJ, Northwest Pacific, were affected with disseminated neoplasia (DN) and confirmed that it was caused by clonally transmitted cancer (CTC), by demonstrating genetic chimerism of mussels with DN and a striking similarity among their “extra” genotypes. The cancer alleles found in our study did not match those of the BTN1 lineage from *M. trossulus* populations on the American coast of the Pacific Ocean but matched the alleles of BTN2 lineage, which has been previously diagnosed in *M. edulis* from Europe and in *M. chilensis* from Chile and Argentine but not in *M. trossulus*, from which it is originally derived from. So, we conclude that *M. trossulus* from the SOJ are infected by BTN2. This finding implies that this species, contrary to the hypothesis of Yonemitsu *et al*.^10^, has not evolved resistance to this disease. Below we will first discuss the pathology and epidemiology of BTN2 in mussels from the SOJ and then its genetic properties.

The features of the neoplastic hemocytes in our study—a rounded shape, a large nucleus, a high nucleus-to-cytoplasm ratio and an increased ploidy—agree with the previous descriptions of DN in *Mytilus*^14,22,23^. What seems unusual is their low proliferation level. Neoplastic hemocytes of mussels are generally assumed to have a high proliferation activity^20^. However, a low proliferation rate of neoplastic hemocytes in mussels with DN was reported in two other studies: of BTN (supposedly BTN2) in France^22^ and of DN^66^ in the same Argentinean population where BTN2 was recognized later^10^. A possible explanation is that in case of BTN2 the proliferation site of the neoplastic cells is located not in the hemolymph. This hypothesis is inspired by the study of Burioli *et al*.^22^, who observed a high mitotic rate of neoplastic cells in the vesicular connective tissue of BTN-infected mussels.

Our data complement the results of another study of DN in mussels from the SOJ^40^, where only one individual with DN was found in a histological examination of 40 mollusks from various localities. We only found DN in one of the three populations examined. The DN prevalence in this Gaydamak Bay population (4.0%) was probably underestimated, being based on flow cytometry, which is not very sensitive at the early stages of the disease^19,22^. Still, our estimate of DN level in the population from the Gaydamak Bay is close to the mean prevalence reported for *Mytilus* populations worldwide^39,67–70^ and much lower than the maximal prevalence of 56% reported for *M. trossulus* population from British Columbia^71^.

Noteworthy, the mussels in the Gaydamak Bay were collected from the surface of a mooring buoy in a heavily polluted area, where no mussels were recorded at the sea floor (our observations). Since BTN is presumably transmitted by cancer cells through the water column (see Caza *et al*.^72^ for more discussion), we suspect that the mussels fouling the buoy contracted the infection from those that had fouled ships moored to it. We point out that mussel populations on mooring buoys, docks etc. may serve as BTN reservoirs while mussel-fouled ships may be the vectors of the disease.

An association between an increased prevalence of DN in mollusks and a high concentration of pollutants, though often surmised, has never been shown explicitly^24^. Our samples from other localities than Gaydamak were limited, and we have no reasons to suppose that mussel populations outside harbors in the SOJ are free from the disease. *M. trossulus* populations in the SOJ and in the Northwest Pacific in general may be a natural reservoir of BTN2, where it is maintained and whence it spreads worldwide.

We recorded four new BTN2 genotypes, in addition to the 11 BTN cancer genotypes reported earlier worldwide (eight of them representing BTN2)^2,10^. We confirm the conclusions of Yonemitsu *et al*.^10^ that BTNs are marked by lineage-specific genotypes at the conservative nuclear EF1α locus and that mitochondrial diversity is observed across BTN2 clones from different individuals as well as within some individuals. This diversity is difficult to interpret at the moment, firstly because of the scarcity of data and secondly because of the possibility of mitochondrial capture from the host and mitochondrial recombination^10,33,73^.

The two main mitochondrial haplotypes defined as the combinations COI-1+CR-1 and COI-2+CR-2, found in homoplasmic condition in our material, were similar but not identical to the two common cancer-associated haplotypes which combine the COI-B alleles with the CR-D and CR-C alleles, respectively, as found by Yonemitsu *et al*.^10^ either in heteroplasmic (Argentina) or homoplasmic state (Europe, CR-D allele).

It is also possible that the apparent homoplasmy of the SOJ mussel cancers is spurious. Yonemitsu *et al*.^10^ have shown using qPCR that the levels of various haplotypes in heteroplasmic cancer clones may be very different. Since we employed molecular cloning of CR and sequenced a limited numbers of colonies (though more than Yonemitsu *et al*.^10^ did to demonstrate heteroplasmy), we might have overlooked rare alleles. Further, if we assume that the cancer clones from Gaydamak are homoplasmic, it follows that the mussels fouling the same mooring buoy were infected with at least two independent clones, which is unlikely, though not impossible.

One novel finding in our study is the remarkable divergence between the two BTN2 haplotypes in the SOJ. Two COI alleles differed from each other and from “B alleles” on average by six substitutions (~1%). This diversity has probably accumulated after the emergence of BTN2. In a study of CTVT mtDNA evolution, Strakova *et al*.^33^ considered three estimates of mutation rate (scaled here to mutations per site per year): based on CTVT nuclear phylogeny (maximum rate estimate 1.23*10^−6^), based on the tempo of accumulation of mutations by human cancers with patient age (1.49*10^−6^) and based on cell divisions adjusted for CTVT generation time (1.09*10^−6^ −5.45*10^−6^). Employing these rates to the 640 bp COI fragment and assuming that each main BTN2 COI lineage acquired on the average three mutations through evolution, we obtain the divergence time estimates of 3885, 3186 and 873-4364 years, respectively. These estimates can hardly be accurate since mitochondrial mutation rates and generation time in DTNs are unknown. A cautious conclusion would be that BTN2 emerged long before DN was first diagnosed in blue mussels in the 1960s^15^.

As to the geographical origin of the BTN2 COI diversity, the comparison with a rich collection of *M. trossulus* COI sequences indicates that it most probably lies outside the Northeast Pacific. Intriguingly, the unique COI allele of the other cancer lineage, BTN1, so far encountered only in British Columbia^10^, also belongs to a haplogroup that is much more common in the Atlantic and the Northwest Pacific than in the Northeast Pacific. Hence our analysis did not unambiguously confirm the endemism of BTN1 to the NE Pacific (Fig. 5).

At least two of the published *M. trossulus* sequences considered in our analyses could indeed turn out to be cancer genotypes. One of them is COI KF931805 from British Columbia^54^, which is identical to the BTN2 allele COI-1. This might mean that BTN2 is cryptically present in British Columbia. If so, DN in mussels from this region could be caused by both BTN1 and BTN2.

Another candidate for a cancerous sequence is KM192133 from the Baltic Sea (genome 62mc10)^37^, which closely matches BTN2 haplotypes (Fig. 4). Yonemitsu *et al*.^10^ considered 62mc10 as a genome of “a normal *M. trossulus* from the Baltic region”, which is close to the ancestral sequence of BTN2, rather than the genome of the cancer itself. There are, however, several reasons to consider the possibility that 62mc10 might be cancerous. Firstly, *M. trossulus* from the Baltic Sea themselves generally lack the typical *M. trossulus* mtDNA. Instead they bear the *M. edulis* mtDNA, as a consequence of introgressive hybridization between these species^74,75^.

In fact, 62mc10 is the only *M. trossulus* like mitochondrial sequence revealed in extensive studies of the Baltic mussels^37^. Secondly, 62mc10 was identified in the mantle (i.e. the gonad) of a female alongside with the standard *M. edulis* F-mtDNA^37^. At the same time, 62mc10, as well as BTN2 mtDNA, is a mosaic sequence: F-mtDNA with M-mtDNA CR segments^10,37^. In mussels, such recombinant mtDNA are usually inherited as standard M-mtDNA (“masculinized” genomes) and so are expected to be found, in a mixture with the F-mtDNA, in gonads of males, not females^52,76^. The mantle tissues are subject to cancer^14^, which can explain the heteroplasmy, and DN has indeed been recorded in the Baltic Sea *M. trossulus*^39^.

The origin of the two cancerous complex mitochondrial haplotypes — so divergent in their control regions, occurring sometimes together and sometimes apart — remains enigmatic. There is little we can add on this matter to the discussion in Yonemitsu *et al*.^10^. BTN2 haplotypes resemble the “masculinized” mitochondrial genomes of mussels. These genomes are common in the Baltic mussels, in which they are *M. edulis*-derived (with the sole exception of 62mc10), and in the hybrid zone between *M. edulis* and *M. trossulus* in Norway, in which they are *M. trossulus*-derived, but rare elsewhere^38,77^. Recombination and masculinization in the Baltic and in Norway are thought to be driven by hybridization between *M. edulis* and *M. trossulus*^37,38^. The ubiquity of masculinized genomes in these two regions prompted Yonemitsu *et al*.^10^ to suggest that BTN2 originated in one of them. If this hypothesis is true, more extensive genetic surveys would find traces of introgression from *M. edulis* in the BTN2 nuclear genome. For example, the absolute majority of the Baltic mussels bear *M. edulis* alleles at the ITS multicopy gene^78^. The lack of *M. edulis* ITS alleles in the hemolymph of cancerous mussels from the SOJ in our study is an additional though not conclusive argument against the Baltic origin of BTN2. While the European origin of BTN2 cannot be refuted, we suggest an alternative hypothesis that it emerged in the heart of *M. trossulus* ancestral range in the Norhwest Pacific.

The similarity between BTN2 and “masculinized” mtDNA might indicate not only the population of the BTN2 origin but also the tissue where it originated and the sex of “patient zero”, the mussel from which BTN2 derive. During embryonic development, paternally inherited mtDNA are maintained in the germline of males^79^, being the genetic marker of male germ cells. This could mean that cancer cells are male germ cells but not hemocytes by origin. If this is true, the first mussel with BTN2 was a male.

## Supporting information

EF1alpha alignment for ML tree

COI alignment for ML tree

CR alignment for ML tree

COI alignment without Crego-Prieto et al, 2015 dataset

COI alignment with Crego-Prieto et al, 2015 dataset

CR alignment for Recombination analysis

## Acknowledgements

We would like to thank Natalia Lentsman for English language editing of the manuscript and Anna Romanovich and Alexey Masharskiy.from the Centre for Molecular and Cell Technologies for technical help. This study was supported by Russian Science Foundation, grant number 19-74-20024.

## Data availability

The sequences generated in this study are deposited in the NCBI GenBank. The accession numbers are listed in Supplementary Table 3. Alignment used for tree generation available as Supplementary files in .fasta format.

## Contributions

MS, NO, and PS designed the study. NO, and MM organized mussel sampling and provided flow cytometry, hemocytology, and immunochemistry analysis. MS and AI provided molecular genetic analysis. MS, RV, and PS carried out the bioinformatics analysis. MS and PS drafted the manuscript. All authors read, approved, and contributed to the final manuscript.

