## Supplementary Figures and Tables for "The root of the transmissible cancer: first description of a widespread *Mytilus trossulus*-derived cancer lineage in *M. trossulus*"

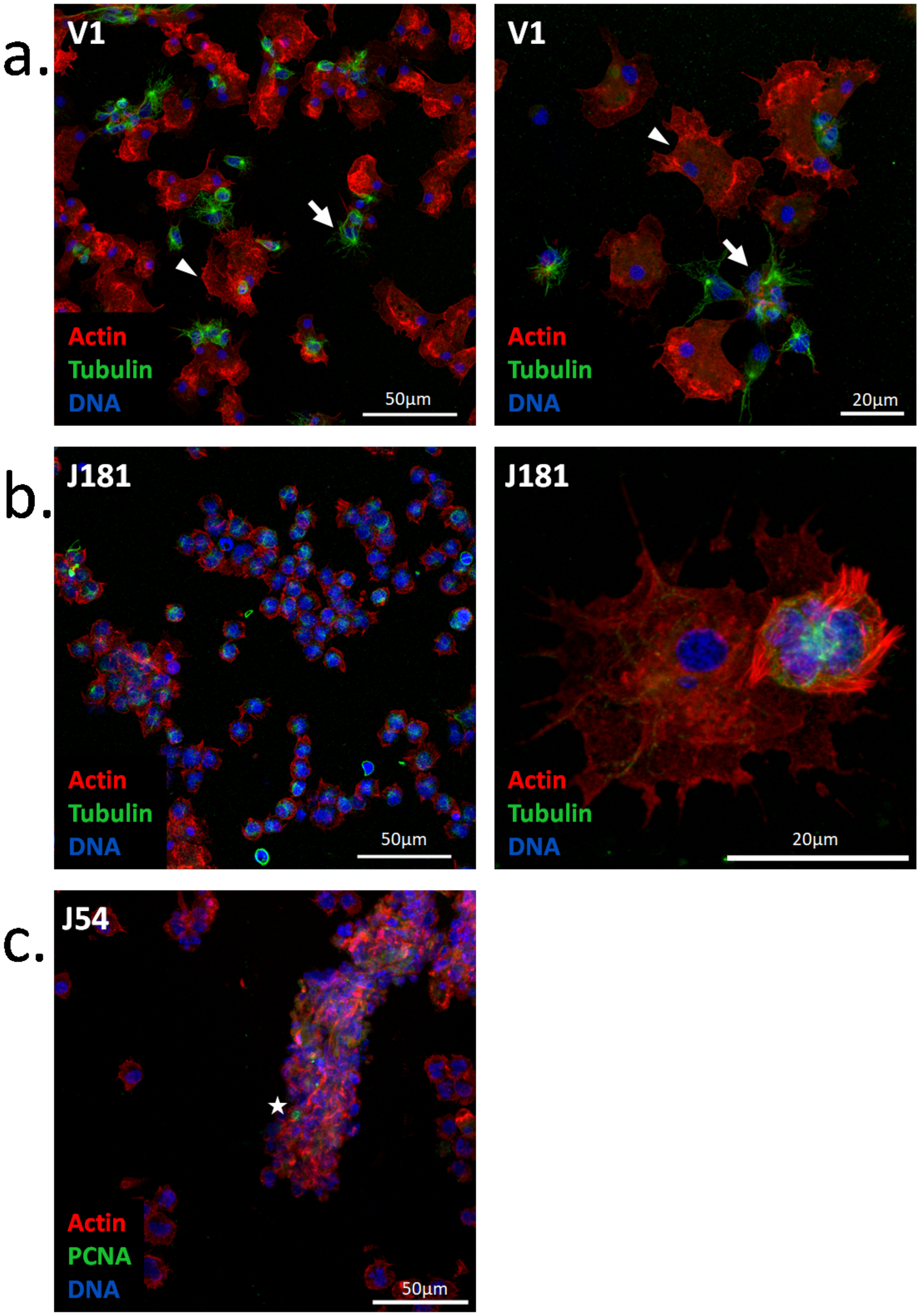


**Figure S1.** *Mytilus trossulus* hemocytes cultivated for 24 hours. Cells were stained with TRITC-labeled phalloidin (red), and the nuclei (chromatin) were stained by DAPI (blue). Hemocytes of a healthy individual V1 (**a**) and a diseased individual J181 (**b**) were stained with tubulin primary antibodies, hemocytes of a diseased individual J54 (**c**) were stained with PCNA antibodies (green). Arrows in pictures of V1 point to actively moving cells, while arrowheads point to adherent cells. Star in picture of J54 marks the cell positive for proliferation.

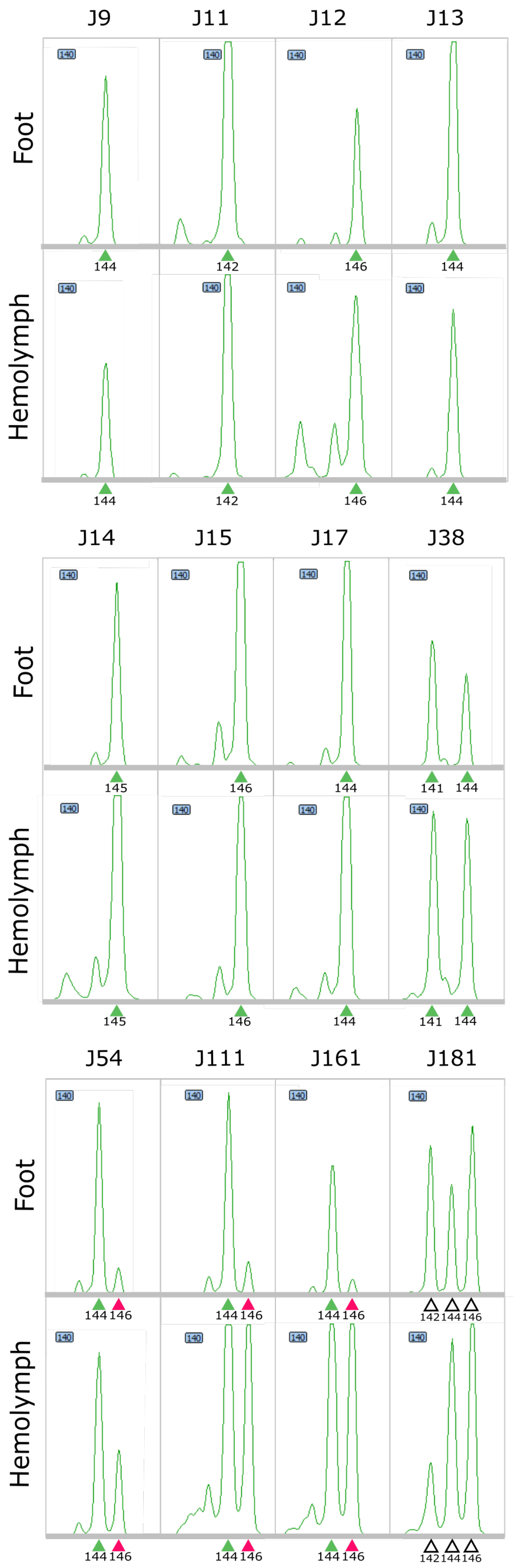


**Figure S2.** Fragment analysis of *Mg*μ3 microsatellite locus from the hemolymph and the foot tissues of eight healthy mussels (control) and four DN-suggested mussels. Colored triangles designate putatively host-derived *Mg*μ3 fragments (green), putatively cancer-derived *Mg*μ3 fragments (violet); open triangles mark unrecognized fragments. The patterns of *Mg*μ3 fragments in the hemolymph and the foot tissues coincide in healthy mussels; an additional peak is present in the hemolymph of DN-suggested mussels.

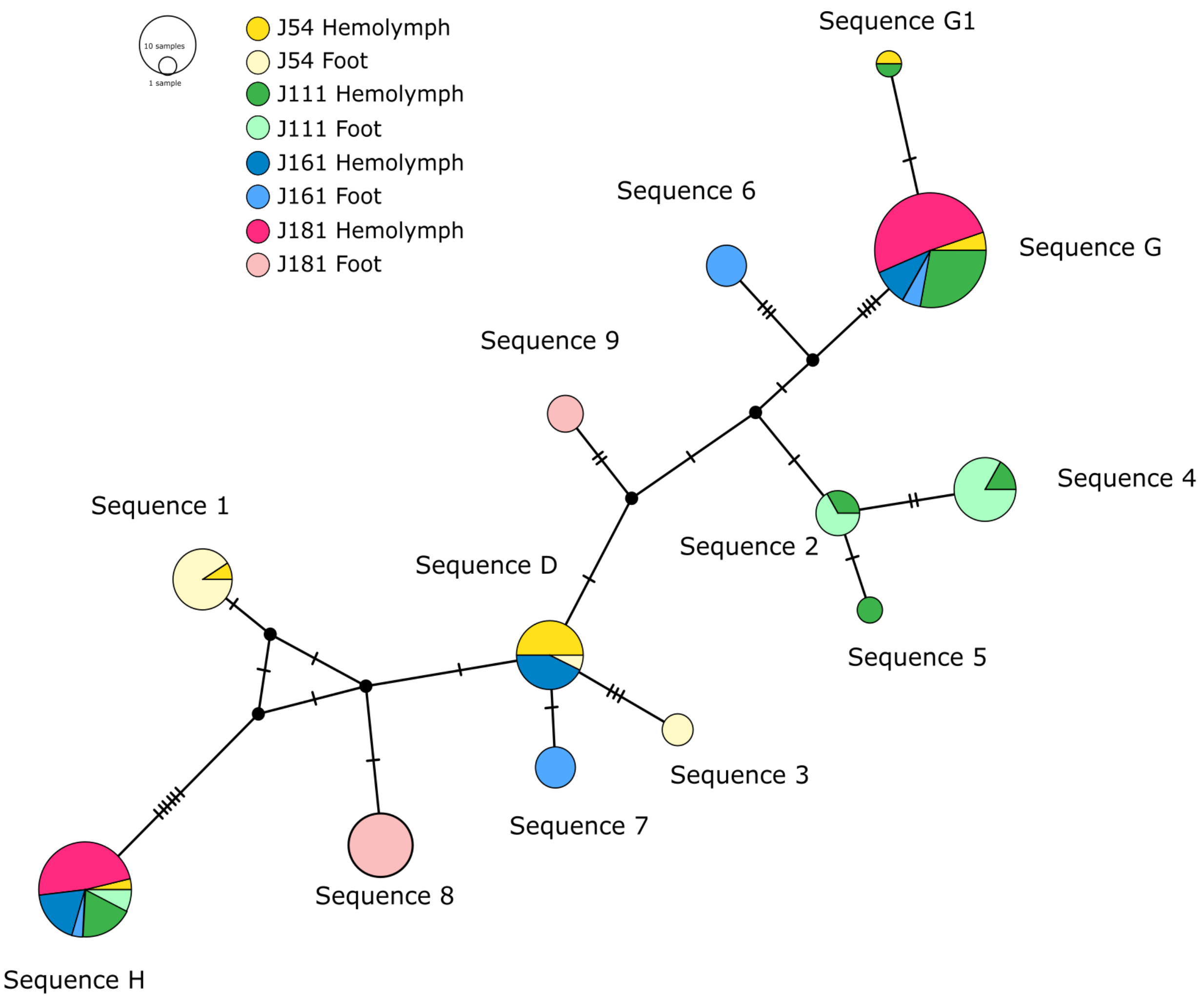


**Figure S3.** TCS network representing EF1α sequences obtained by molecular cloning. Sequences from individual mussels and from different tissues (hemolymph and foot) are color-coded (see legend on the top). All minor sequences represented by a single bacterial colony have been removed.

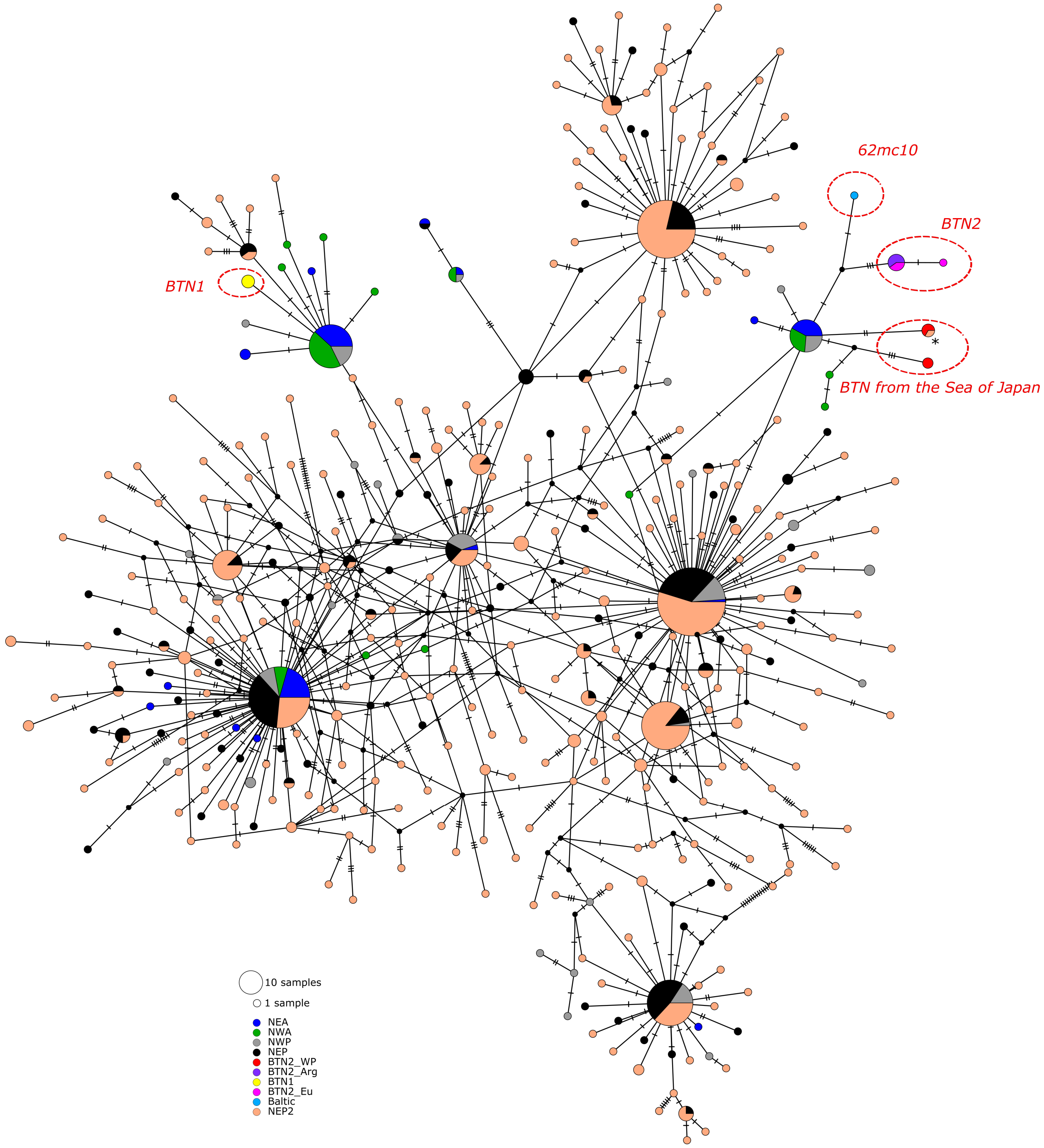


**Figure S4.** *M. trossulus* and BTN COI haplotype network from the TCS analysis of 542-bp alignment of 843 sequences. The data come from a reanalysis of the results (Fig. 5) with additional 493 sequences from Crego-Prieto *et al.*[54]. Each circle represents a single allele. The size of the circle is proportional to the number of individuals found to bear the allele. Bars indicate mutations between alleles. Small black circles indicate hypothetical haplotypes predicted by the model. The alignment is available as Supplementary Data S5. The geographical origin of samples is color-coded. Samples corresponding to BTN1 (British Columbia) are in yellow, to BTN2 in pink (Europe, BTN2_Eu) and purple (Argentina, BTN2_Arg), to BTN from the SOJ in red (BTN2_WP). Reference *M. trossulus* samples from the Northwest Pacific (NWP) are in grey, from Northeast Pacific (NEP), in black, from Northwest Atlantic (NWA), in green, from Northeast Atlantic (NEA), in dark blue, and from the Baltic Sea (a single sample 62mc10), in light blue (Baltic). The data from Northeast Pacific from Crego-Prieto et al.[54] are in orange

(NEP2). The asterisk mark the putatively cancer sequence (NCBI accession number KF931805) from Crego-Prieto et al.[54]

**a.**

| | D alleles, p-value | C alleles, p-value |
|---|---|---|
| RDP | - | - |
| GENECONV | $4.668*10^{-5}$ | - |
| Bootscan | $5.704*10^{-3}$ | - |
| Maxchi | $1.556*10^{-9}$ | $4.229*10^{-2}$ |
| Chimaera | $1.145*10^{-4}$ | - |
| SiScan | $6.138*10^{-11}$ | $1.248*10^{-17}$ |
| 3Seq | $1.042*10^{-14}$ | $9.983*10^{-6}$ |

**b.**

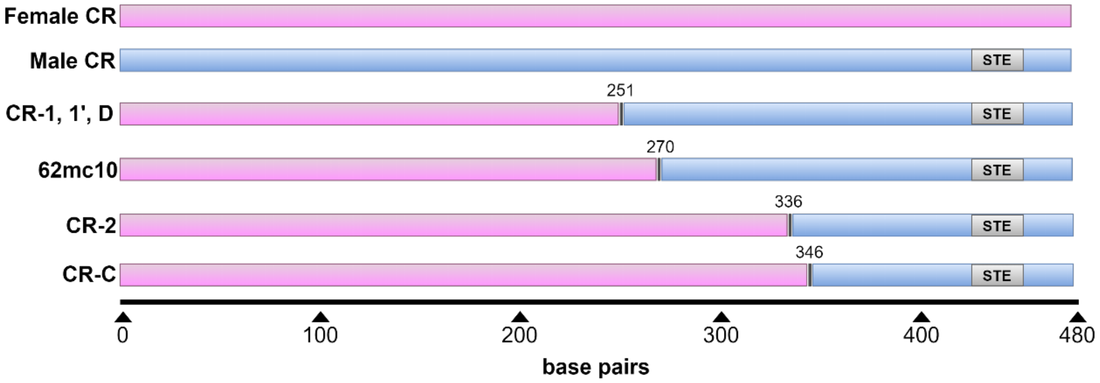


**Figure S5.** (**a**) 479 bp long alignment subjected to recombination essay in RDP4 package with default parameters included CR alleles of cancerous mussels detected in this study and in the study of Yonemitsu *et al*. (2019), CR of 62mc10 individual from the Baltic Sea and the set of reference male and female CR (see Accession numbers in the alignment included as Supplementary data S6). The RDP4 package detected two recombination events in the region around 249-263 nucleotide position of alignment in CR-D, D1, D2, D3, 1, 1' and 62mc10 and in the region of 333-360 nucleotide position in CR-C, C1 and 2. RDP4 output p-values (cutoff of 0.05) are represented in the table. (**b**) The putative breakpoints marked at schematically depicted alignment of CR used in the recombination analysis. Each bar represents a sequence. Pink color designates standard F-mtDNA, blue color, standard M-mtDNA. The putative breakpoints are marked with black vertical lines, designated by the nucleotide position from the start of alignment. Sperm transmissible element is marked with a grey box.

### Supplementary tables

**Table S1.** The list of *Mytilus trossulus* and BTN COI sequences used for creating TCS haplotype networks.

| NCBI Accession numbers | N of sequences | Source |
|---|---|---|
| AY823625 | 1 | Breton et al., 2006 |
| KF643248, KF643275, KF643297, KF643331, KF643385, KF643387, KF643419, KF643426, KF643427, KF643460, KF643487, KF643497, KF643502, KF643600, KF643628, KF643676, KF643701, KF643735, KF643750, KF643757, KF643770, KF643789, KF643795, KF643812, KF643820, KF643889, KF643915, KF643952, KF643955, KF643956, KF644043, KF644059, KF644073, KF644107, KF644123, KF644190, KF644206, KF644232, KF644263, KF644321, KF644327 | 41 | Layton et al., 2014 |
| GQ902548-GQ902685 | 138 | Marko et al., 2010 |
| MN546832-MN546835, MN546842, MN546848, MN546852-MN546858 | 15 | Yonemitsu et al., 2019 |
| KM192133 | 1 | Zbawicka et al., 2014 |
| MN119673 | 1 | Chung et al., 2019 |
| MG422062 | 1 | deWaard et al., 2019 |
| MT736560-MT736683 | 124 | Laakkonen et al., 2020 |
| KF931763-KF931851, KF931854-KF931898, KF931901-KF931939, KF931943-KF932262 | 493 | Crego-Prieto et al., 2015 |
| MW191702-MW191719, MT857958-MT857963, MW150800-MW150801 | 28 | This study |

**Table S2.** The list of EF1α and CR sequences revealed by molecular cloning.

| Locus | Individual | Tissue | Clone | Haplotype |
|---|---|---|---|---|
| EF1α | J17 | Hemolymph | 2 | D |
| EF1α | J17 | Hemolymph | 3 | D |
| EF1α | J17 | Hemolymph | 5 | D |
| EF1α | J17 | Hemolymph | 8 | D |
| EF1α | J17 | Hemolymph | 9 | D' |
| EF1α | J17 | Hemolymph | 10 | D' |
| EF1α | J17 | Hemolymph | 18 | D' |
| EF1α | J17 | Hemolymph | 24 | D' |
| EF1α | J17 | Hemolymph | 26 | D' |
| EF1α | J17 | Hemolymph | 27 | D' |
| EF1α | J17 | Hemolymph | 30 | D' |
| EF1α | J17 | Hemolymph | 32 | D |
| EF1α | J17 | Hemolymph | 33 | D' |
| EF1α | J17 | Hemolymph | 34 | D' |
| EF1α | J17 | Hemolymph | 35 | D' |
| EF1α | J17 | Hemolymph | 36 | D |
| EF1α | J38 | Hemolymph | 1 | 10 |
| EF1α | J38 | Hemolymph | 2 | 10 |
| EF1α | J38 | Hemolymph | 3 | 10 |
| EF1α | J38 | Hemolymph | 4 | 10 |
| EF1α | J38 | Hemolymph | 6 | 10 |
| EF1α | J38 | Hemolymph | 9 | 10 |
| EF1α | J38 | Hemolymph | 10 | 10 |
| EF1α | J38 | Hemolymph | 11 | 10 |
| EF1α | J38 | Hemolymph | 13 | 10 |
| EF1α | J38 | Hemolymph | 15 | 10 |
| EF1α | J38 | Hemolymph | 16 | 10 |
| EF1α | J38 | Hemolymph | 17 | 10 |
| EF1α | J38 | Hemolymph | 18 | 10 |
| EF1α | J38 | Hemolymph | 19 | 10 |
| EF1α | J38 | Hemolymph | 21 | 10 |
| EF1α | J38 | Hemolymph | 27 | 10 |
| EF1α | J54 | Hemolymph | 1 | MINOR |
| EF1α | J54 | Hemolymph | 2 | D |
| EF1α | J54 | Hemolymph | 5 | MINOR |
| EF1α | J54 | Hemolymph | 6 | H |
| EF1α | J54 | Hemolymph | 9 | MINOR |
| EF1α | J54 | Hemolymph | 11 | MINOR |
| EF1α | J54 | Hemolymph | 15 | MINOR |
| EF1α | J54 | Hemolymph | 16 | D |
| EF1α | J54 | Hemolymph | 17 | D |
| EF1α | J54 | Hemolymph | 18 | MINOR |
| EF1α | J54 | Hemolymph | 19 | D |
| EF1α | J54 | Hemolymph | 20 | D |
| EF1α | J54 | Hemolymph | 21 | D |
| EF1α | J54 | Hemolymph | 24 | MINOR |
| EF1α | J54 | Hemolymph | 26 | D |
| EF1α | J54 | Hemolymph | 31 | G1 |
| EF1α | J54 | Hemolymph | 33 | G |
| EF1α | J54 | Hemolymph | 41 | 1 |

| | | | | |
|---|---|---|---|---|
| EF1α | J54 | Hemolymph | 49 | G |
| EF1α | J54 | Foot | 3 | 1 |
| EF1α | J54 | Foot | 4 | 1 |
| EF1α | J54 | Foot | 5 | 1 |
| EF1α | J54 | Foot | 6 | 1 |
| EF1α | J54 | Foot | 7 | MINOR |
| EF1α | J54 | Foot | 8 | 1 |
| EF1α | J54 | Foot | 9 | D |
| EF1α | J54 | Foot | 10 | 3 |
| EF1α | J54 | Foot | 11 | 3 |
| EF1α | J54 | Foot | 12 | 1 |
| EF1α | J54 | Foot | 13 | 1 |
| EF1α | J54 | Foot | 14 | 1 |
| EF1α | J54 | Foot | 15 | 3 |
| EF1α | J54 | Foot | 19 | 1 |
| EF1α | J54 | Foot | 17 | 1 |
| EF1α | J54 | Foot | 18 | MINOR |
| EF1α | J111 | Hemolymph | 2 | G |
| EF1α | J111 | Hemolymph | 3 | G |
| EF1α | J111 | Hemolymph | 7 | G |
| EF1α | J111 | Hemolymph | 9 | G |
| EF1α | J111 | Hemolymph | 11 | MINOR |
| EF1α | J111 | Hemolymph | 12 | 5 |
| EF1α | J111 | Hemolymph | 13 | G |
| EF1α | J111 | Hemolymph | 14 | H |
| EF1α | J111 | Hemolymph | 15 | 5 |
| EF1α | J111 | Hemolymph | 17 | H |
| EF1α | J111 | Hemolymph | 18 | G |
| EF1α | J111 | Hemolymph | 19 | 4 |
| EF1α | J111 | Hemolymph | 20 | G |
| EF1α | J111 | Hemolymph | 24 | G |
| EF1α | J111 | Hemolymph | 25 | H |
| EF1α | J111 | Hemolymph | 27 | G |
| EF1α | J111 | Hemolymph | 28 | G |
| EF1α | J111 | Hemolymph | 29 | 2 |
| EF1α | J111 | Hemolymph | 30 | H |
| EF1α | J111 | Hemolymph | 31 | G1 |
| EF1α | J111 | Hemolymph | 32 | 2 |
| EF1α | J111 | Hemolymph | 33 | G |
| EF1α | J111 | Hemolymph | 34 | MINOR |
| EF1α | J111 | Hemolymph | 35 | 4 |
| EF1α | J111 | Hemolymph | 37 | H |
| EF1α | J111 | Foot | 3 | 4 |
| EF1α | J111 | Foot | 4 | 4 |
| EF1α | J111 | Foot | 5 | 4 |
| EF1α | J111 | Foot | 9 | 4 |
| EF1α | J111 | Foot | 14 | 4 |
| EF1α | J111 | Foot | 16 | 4 |
| EF1α | J111 | Foot | 17 | 4 |
| EF1α | J111 | Foot | 18 | H |
| EF1α | J111 | Foot | 19 | 2 |
| EF1α | J111 | Foot | 20 | 2 |
| EF1α | J111 | Foot | 25 | 4 |
| EF1α | J111 | Foot | 27 | 2 |
| EF1α | J111 | Foot | 33 | H |

| | | | | |
|---|---|---|---|---|
| EF1α | J111 | Foot | 34 | 4 |
| EF1α | J111 | Foot | 37 | 4 |
| EF1α | J111 | Foot | 38 | 2 |
| EF1α | J161 | Hemolymph | 1 | H |
| EF1α | J161 | Hemolymph | 3 | G |
| EF1α | J161 | Hemolymph | 6 | D |
| EF1α | J161 | Hemolymph | 7 | D |
| EF1α | J161 | Hemolymph | 9 | G |
| EF1α | J161 | Hemolymph | 10 | G |
| EF1α | J161 | Hemolymph | 13 | H |
| EF1α | J161 | Hemolymph | 15 | G |
| EF1α | J161 | Hemolymph | 18 | MINOR |
| EF1α | J161 | Hemolymph | 20 | G |
| EF1α | J161 | Hemolymph | 21 | MINOR |
| EF1α | J161 | Hemolymph | 22 | D |
| EF1α | J161 | Hemolymph | 23 | G |
| EF1α | J161 | Hemolymph | 24 | MINOR |
| EF1α | J161 | Hemolymph | 25 | D |
| EF1α | J161 | Hemolymph | 26 | D |
| EF1α | J161 | Hemolymph | 27 | G |
| EF1α | J161 | Hemolymph | 28 | MINOR |
| EF1α | J161 | Hemolymph | 29 | MINOR |
| EF1α | J161 | Hemolymph | 30 | H |
| EF1α | J161 | Hemolymph | 31 | H |
| EF1α | J161 | Hemolymph | 33 | MINOR |
| EF1α | J161 | Hemolymph | 34 | D |
| EF1α | J161 | Hemolymph | 35 | H |
| EF1α | J161 | Foot | 1 | MINOR |
| EF1α | J161 | Foot | 2 | MINOR |
| EF1α | J161 | Foot | 5 | 7 |
| EF1α | J161 | Foot | 6 | G |
| EF1α | J161 | Foot | 7 | 6 |
| EF1α | J161 | Foot | 10 | G |
| EF1α | J161 | Foot | 11 | 7 |
| EF1α | J161 | Foot | 14 | MINOR |
| EF1α | J161 | Foot | 18 | 7 |
| EF1α | J161 | Foot | 20 | H |
| EF1α | J161 | Foot | 21 | 6 |
| EF1α | J161 | Foot | 23 | 6 |
| EF1α | J161 | Foot | 28 | 6 |
| EF1α | J161 | Foot | 33 | 7 |
| EF1α | J161 | Foot | 35 | MINOR |
| EF1α | J161 | Foot | 36 | 6 |
| EF1α | J181 | Hemolymph | 1_pJet | G |
| EF1α | J181 | Hemolymph | 2_pJet | G |
| EF1α | J181 | Hemolymph | 3_pJet | G |
| EF1α | J181 | Hemolymph | 4_pJet | G |
| EF1α | J181 | Hemolymph | 5_pJet | G |
| EF1α | J181 | Hemolymph | 6_pJet | G |
| EF1α | J181 | Hemolymph | 7_pJet | G |
| EF1α | J181 | Hemolymph | 8_pJet | MINOR |
| EF1α | J181 | Hemolymph | 9_pJet | H |
| EF1α | J181 | Hemolymph | 10_pJet | G |
| EF1α | J181 | Hemolymph | 11_pJet | H |
| EF1α | J181 | Hemolymph | 12_pJet | G |

| | | | | |
|---|---|---|---|---|
| EF1α | J181 | Hemolymph | 13_pJet | G |
| EF1α | J181 | Hemolymph | 14_pJet | G |
| EF1α | J181 | Hemolymph | 15_pJet | H |
| EF1α | J181 | Hemolymph | 16_pJet | MINOR |
| EF1α | J181 | Hemolymph | 1 | MINOR |
| EF1α | J181 | Hemolymph | 2 | G |
| EF1α | J181 | Hemolymph | 5 | MINOR |
| EF1α | J181 | Hemolymph | 6 | H |
| EF1α | J181 | Hemolymph | 7 | H |
| EF1α | J181 | Hemolymph | 8 | G |
| EF1α | J181 | Hemolymph | 9 | H |
| EF1α | J181 | Hemolymph | 10 | G |
| EF1α | J181 | Hemolymph | 12 | G |
| EF1α | J181 | Hemolymph | 13 | G |
| EF1α | J181 | Hemolymph | 15 | G |
| EF1α | J181 | Hemolymph | 16 | G |
| EF1α | J181 | Hemolymph | 18 | H |
| EF1α | J181 | Hemolymph | 22 | G |
| EF1α | J181 | Hemolymph | 23 | H |
| EF1α | J181 | Hemolymph | 24 | G |
| EF1α | J181 | Hemolymph | 25 | H |
| EF1α | J181 | Hemolymph | 26 | H |
| EF1α | J181 | Hemolymph | 28 | H |
| EF1α | J181 | Hemolymph | 29 | H |
| EF1α | J181 | Hemolymph | 30 | MINOR |
| EF1α | J181 | Hemolymph | 31 | G |
| EF1α | J181 | Hemolymph | 32 | G |
| EF1α | J181 | Hemolymph | 33 | G |
| EF1α | J181 | Hemolymph | 34 | MINOR |
| EF1α | J181 | Hemolymph | 35 | G |
| EF1α | J181 | Hemolymph | 36 | H |
| EF1α | J181 | Foot | 2 | 8 |
| EF1α | J181 | Foot | 4 | 8 |
| EF1α | J181 | Foot | 6 | 8 |
| EF1α | J181 | Foot | 16 | 8 |
| EF1α | J181 | Foot | 17 | 9 |
| EF1α | J181 | Foot | 20 | 9 |
| EF1α | J181 | Foot | 23 | 8 |
| EF1α | J181 | Foot | 24 | 9 |
| EF1α | J181 | Foot | 28 | 8 |
| EF1α | J181 | Foot | 29 | 8 |
| EF1α | J181 | Foot | 30 | 8 |
| EF1α | J181 | Foot | 31 | 8 |
| EF1α | J181 | Foot | 39 | 9 |
| EF1α | J181 | Foot | 47 | 8 |
| EF1α | J181 | Foot | 51 | 8 |
| EF1α | J181 | Foot | 58 | 8 |
| CR | J54 | Hemolymph | 1 | 1 |
| CR | J54 | Hemolymph | 2 | 1 |
| CR | J54 | Hemolymph | 4 | 1 |
| CR | J54 | Hemolymph | 5 | 1 |
| CR | J54 | Hemolymph | 6 | 1 |
| CR | J54 | Hemolymph | 7 | 1 |
| CR | J54 | Hemolymph | 8 | 1 |
| CR | J54 | Hemolymph | 10 | 1 |

| | | | | |
|---|---|---|---|---|
| CR | J54 | Hemolymph | 11 | 1 |
| CR | J54 | Hemolymph | 12 | 1 |
| CR | J54 | Hemolymph | 14 | 1 |
| CR | J54 | Hemolymph | 15 | 1 |
| CR | J54 | Hemolymph | 16 | 1 |
| CR | J54 | Hemolymph | 19 | 1 |
| CR | J54 | Hemolymph | 20 | 1 |
| CR | J54 | Hemolymph | 21 | 1 |
| CR | J111 | Hemolymph | 1 | 2 |
| CR | J111 | Hemolymph | 3 | 2 |
| CR | J111 | Hemolymph | 6 | 2 |
| CR | J111 | Hemolymph | 12 | 2 |
| CR | J111 | Hemolymph | 15 | 2 |
| CR | J111 | Hemolymph | 2 | 4 |
| CR | J111 | Hemolymph | 7 | 4 |
| CR | J111 | Hemolymph | 10 | 4 |
| CR | J111 | Hemolymph | 13 | 4 |
| CR | J111 | Hemolymph | 14 | 4 |
| CR | J111 | Hemolymph | 16 | 4 |
| CR | J111 | Hemolymph | 8 | MINOR |
| CR | J111 | Hemolymph | 4 | MINOR |
| CR | J111 | Hemolymph | 11 | MINOR |
| CR | J111 | Hemolymph | 17 | MINOR |
| CR | J161 | Hemolymph | 1 | 1 |
| CR | J161 | Hemolymph | 2 | 1' |
| CR | J161 | Hemolymph | 3 | 1' |
| CR | J161 | Hemolymph | 4 | 1 |
| CR | J161 | Hemolymph | 5 | 1 |
| CR | J161 | Hemolymph | 6 | 1 |
| CR | J161 | Hemolymph | 7 | 1 |
| CR | J161 | Hemolymph | 8 | 1 |
| CR | J161 | Hemolymph | 9 | 1' |
| CR | J161 | Hemolymph | 10 | 1 |
| CR | J161 | Hemolymph | 11 | 1' |
| CR | J161 | Hemolymph | 13 | 1 |
| CR | J161 | Hemolymph | 14 | 1' |
| CR | J161 | Hemolymph | 15 | 1 |
| CR | J161 | Hemolymph | 16 | 1 |
| CR | J161 | Hemolymph | 17 | MINOR |
| CR | J181 | Hemolymph | 1 | 2 |
| CR | J181 | Hemolymph | 5 | 2 |
| CR | J181 | Hemolymph | 6 | 2 |
| CR | J181 | Hemolymph | 10 | 2 |
| CR | J181 | Hemolymph | 11 | 2 |
| CR | J181 | Hemolymph | 12 | 2 |
| CR | J181 | Hemolymph | 15 | 2 |
| CR | J181 | Hemolymph | 16 | 2 |
| CR | J181 | Hemolymph | 17 | 2 |
| CR | J181 | Hemolymph | 4 | 6 |
| CR | J181 | Hemolymph | 7 | 6 |
| CR | J181 | Hemolymph | 9 | 6 |
| CR | J181 | Hemolymph | 2 | MINOR |
| CR | J181 | Hemolymph | 13 | MINOR |
| CR | J181 | Hemolymph | 20 | MINOR |
| CR | J181 | Hemolymph | 14 | MINOR |

**Table S3.** NCBI accession numbers of cancer and host sequences generated in this study.

| Sequence | Suggested origin | NCBI Accession Number |
|---|---|---|
| COI-1 | Cancer | MT857958 |
| COI-2 | Cancer | MT857959 |
| COI-3 | Host | MT857960 |
| COI-4 | Host | MT857961 |
| COI-5 | Host | MT857962 |
| COI-6 | Host | MT857963 |
| COI-7 | Host | MW150800 |
| COI-8 | Host | MW150801 |
| CR-1 | Cancer | MT877229 |
| CR-1' | Cancer | MT877228 |
| CR-2 | Cancer | MT877230 |
| CR-3 | Host | MT877231 |
| CR-4 | Host | MT877233 |
| CR-5 | Host | MT877232 |
| CR-6 | Host | MT877234 |
| CR-7 | Host | MW013817 |
| CR-8 | Host | MW013818 |
| EF1α-1 | Host | MW187821 |
| EF1α-2 | Host | MW187822 |
| EF1α-3 | Host | MW187823 |
| EF1α-4 | Host | MW187824 |
| EF1α-5 | Host | MW187825 |
| EF1α-6 | Host | MW187826 |
| EF1α-7 | Host | MW187827 |
| EF1α-8 | Host | MW187828 |
| EF1α-9 | Host | MW187829 |
| EF1α-10 | Host | MW187830 |
| EF1α-D | Host | MW187831 |
| EF1α-D' | Host | MW187832 |
| EF1α-G1 | Cancer | MW187833 |
| EF1α-H | Cancer | MW187834 |
| EF1α-G | Cancer | MW187835 |
| COI-R7 | Host | MW191702 |
| COI-R11 | Host | MW191703 |
| COI-R13 | Host | MW191704 |
| COI-R14 | Host | MW191705 |
| COI-J10 | Host | MW191706 |
| COI-J11 | Host | MW191707 |
| COI-J12 | Host | MW191708 |
| COI-J13 | Host | MW191709 |
| COI-J14 | Host | MW191710 |
| COI-J15 | Host | MW191711 |
| COI-J16 | Host | MW191712 |
| COI-J2 | Host | MW191713 |
| COI-J3 | Host | MW191714 |
| COI-J4 | Host | MW191715 |
| COI-J6 | Host | MW191716 |
| COI-J7 | Host | MW191717 |
| COI-J8 | Host | MW191718 |
| COI-J9 | Host | MW191719 |
